# How human intervention and climate change shaped the fate of the Northern Bald Ibis from ancient Egypt to the presence: an interdisciplinary approach to extinction and recovery of an iconic bird species

**DOI:** 10.1101/2020.11.25.397570

**Authors:** Johannes Fritz, Jiří Janák

## Abstract

Once widespread around the Mediterranean, the Northern Bald Ibis (*Geronticus eremita*) became one of the rarest birds in the world. We trace the history of this species through different epochs to the present. A particular focus is on its life and disappearance in ancient Egypt, where it attained the greatest mythological significance as a hieroglyphic sign for ‘blessed ancestor spirits’, and on modern endeavours to rewild and restore the species. The close association of the Northern Bald Ibis with human culture in ancient Egypt, as in other regions, is caused by primarily two reasons, the characteristic appearance and behaviour, as well as the need for open foraging areas. In consequence, a mutualistic relationship between humans and birds was formed in some cultures. The benefit for the Northern Bald Ibis was mainly the availability of feeding habitats, which were cleared by humans for farming or grazing and might have contributed to the spread of the species. The benefit to people was primarily cultural and mythological, whereby the bird was worshiped in ancient Egypt and in Muslim cultures, while Christian cultures in Europe rather regarded it as bad omen or nuisance, like any black bird species. Another benefit was profane in nature, the species was also hunted for food, mainly in Europe. But alike many other species, proximity to humans also carried a high risk for the Northern Bald Ibis. We discuss various kinds of human impacts that were driving causes for the extinction of the species in almost all regions. However, the historical disappearance of populations also correlates markedly with changes in climate, especially in ancient Egypt and the Middle Ages. This fact has important implications for current conservation efforts, especially since international action plans for the Northern Bald Ibis have taken little account of climate change effects so far. The Northern Bald Ibis is an outstanding example of how an interdisciplinary cultural-historical and natural-scientific approach significantly promotes the interpretation of historical evidence as well as the implementation of current rewilding and restoration efforts.

## Introduction

There is a growing consensus that the rapidly increasing challenges of biodiversity conservation and restoration cannot be overcome without a thorough knowledge of human society and its history (1). In an interdisciplinary approach social science can promote the sustainability and effectiveness of conservation measures, if certain obstacles can be overcome, like suitable platforms for interdisciplinary publications or misunderstandings due to different perspectives or definitions (2). On the other hand side, cooperation with natural scientists can also drive research in the social sciences, such as the interpretation of historical events and depictions (3).

The Northern Bald Ibis (*Geronticus eremita*) combines current conservation needs and cultural-historical research questions in an extraordinary way. It is among the most threatened bird species in the world, listed on the IUCN Red List as *critically endangered* for 24 year, before it was downgraded to *endangered* in 2018 due to extensive protection efforts (4). The unique appearance, conspicuous behaviour and diverse flight characteristics made this migratory bird species a noticeable creature on the ground and in the air. Moreover, humans created habitats for this species when they cultivated cropland and meadows in different regions and epochs. That inevitably made him a species whose history and fate were closely interwoven with that of the human cultures with which he interacted and shared the habitats (5–7).

An epoch with particularly rich historical traces of the Northern Bald Ibis was the ancient Egypt (8). A depiction of this species was used in Egyptian scripts as the *akh*-sigh, which is easily recognizable characteristic shape of the bird’s body, with the long-curved bill and most characteristic the typical crest covering the back. As a hieroglyphic sign for ‘blessed ancestor spirits’ it attained the greatest mythological significance. However, there are no attestations of keeping, hunting or sacrificing or mummifying the Northern Bald Ibis, as it was the case with the Sacred Ibis (*Threskiornis aethiopicus*)or the Glossy Ibis (*Plegadis falcinellus*).

In this article we aim to shed light on the peculiarities in the historic context and the circumstance of extinction of the species in the ancient Egypt and in other periods and regions, using current knowledge on the biology of this species. We also want to draw conclusions from this that are relevant to the current and urgently needed conservation measures for the species.

## Material and Methods

Studies on Northern Bald Ibis behaviour were mainly be done in the frame of the European LIFE+ reintroduction project (LIFE+12-BIO_AT_000143; www.waldrapp.eu). This project aims to establish a migratory population in Central Europe with a migration tradition to the southern Tuscany. A steadily increasing number of wild living birds migrate between the common wintering site and three breeding sites north of the Alps. Release in course of this project is done in accordance with the IUCN Reintroduction Guidelines and is approved by the regarding national authorities.

Studies on the Egyptian concept of the *akh* (represented in hieroglyphic script by a depiction of the Northern Bald Ibis) have been undertaken for more than 10 years within several broader research projects of the Czech Institute of Egyptology, Charles University, Prague. Methodologically, this research covered archaeological excavation, analysis and interpretation of material, textual and iconographical evidence, analysis of all available original textual sources dealing with the ancient Egyptian religious concept of the *akh*, collecting (and interpreting) evidence on bodily remains of the Northern Bald Ibis from Egypt and, most importantly, building a palaeographic database of the *akh* depictions from all historic periods of ancient Egypt. Thus, we were able to gain different kinds of data (material, as well as pictorial and textual) on the presence and absence of the Northern Bald Ibis in ancient Egypt.

## Results

### The Northern Bald Ibis

Once a widespread colonial migratory bird species, the Northern Bald Ibis (*Geronticus eremita*; Fig 1) had to be put on the Red List in 1994 as one of the rarest birds in the world. At that time, only one population remained in the wild, living at the Atlantic coast in Morocco. But also this population was at a critical population size with only 65 breeding pairs (9). They were the remainders of a once widespread species with breeding colonies on the African continent from Morocco to Egypt, in large parts of Europe and on the Arabian Peninsula (10).

**Fig 1.**
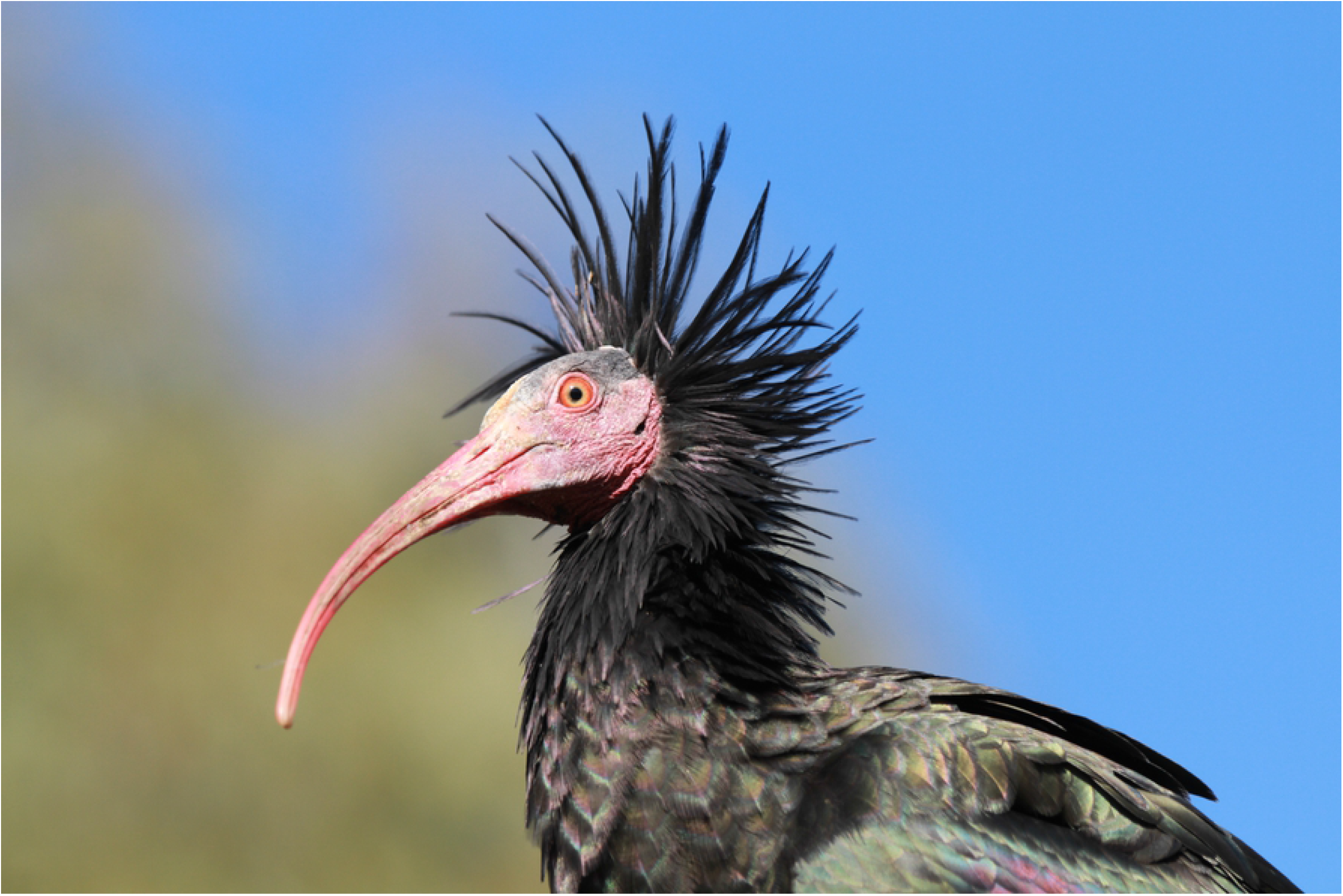
Portrait of an adult Northern Bald Ibis. Picture J Fritz.

The Europe, the species went extinct already in the Middle Age. From the publications of the Swiss naturalist Conrad Gesner (11) and other sources we are able to reconstruct historic breeding sites mainly along the northern foothills of the Alps (12–14). But there is also increasing evidence for a larger former breeding range in Europe, with indications for southern Spain (15), the Upper Adriatic Region (16), Bulgaria (17) or the *Kaiserstuhl* region in Baden-Württemberg with bones dating to the 4^th^ century AD (18). Northern Bald Ibis bones found in a cave located in the French department of Ardèche even dated to the Iron Age, between 764 and 406 BC (19).

The historic evidence for a long-lasting presence of the Northern Bald Ibis in Europe throughout the Holocene corresponds with recent genetic findings. In an extensive analysis of mitochondrial DNA, Wirtz et al. (2018) found no genetic differentiation between the Moroccan population and the former Middle East population. This indicates a contiguous Northern Bald Ibis population whose breeding range covered large parts of North Africa, Europe and the Middle East.

The separation into a western and eastern populations took place when the species disappeared from Europe in the early 17^th^ century. Historic record clearly indicated anthropogenic causes, mainly hunting and collection of chicks (21). But the rapid decline in Europe was probably accelerated by the Little Ice Age, as it is also evident for other species (22). The period from the beginning of the worldwide glacial expansion in 1550 till the first climatic minimum in 1650 fits very well with the decline of the Northern Bald Ibis population. The deteriorating climatic condition may have led to reduced breeding success and increased loss rate due to onset of winter (16,23).

After vanishing in Europe, the cultural memory of the species was lost for centuries and historic depictions were taken for portraits of a mythical creature. It was not until 1897 that ornithologists recognized that the depictions resemble a real living species which was described for the Middle East and only then did the species receive its current scientific name *Geronticus eremita* (13).

At the time when the species disappeared in Europe, it was still widespread in the Middle East with some colonies holding several thousand individuals (24). However, till end of the 20^th^ century all these colonies became exterminated. Major causes were destruction of the habitat, disturbance of the breeding colonies and the industrialization of agriculture. A well-documented example is the former colony along the river Euphrates, near the town of Bireçik in southern Turkey. There, the intensive application of DDT against malaria and locusts caused the loss of more than 600 individuals, about 70% of the population, between 1959 and 1960 (25,26). In 1989, from three remaining adult birds, which returned from migration to this breeding site, only one bird survived to the end of the breeding season.

This was generally assumed to be the end of the last wild colony in the Middle East (27). But in fact, it was not. Very unexpectedly, a small relict population was discovered in 2002 near Palmyra in Syria, comprising of only seven individuals (28). Satellite tracking revealed that they still migrate over more than 3000 km to the historic wintering site near Addis Ababa in Ethiopia (29). The same birds which behaved very shy at their breeding site in the Syrian desert lived during winter in an agro-pastoral landscape in the close surrounding of villages in Ethiopia (30).

After the surprising discovery of this relic population, extensive international conservation efforts followed. They even included the release of three juveniles from a semi-captive breeding colony in Bireçik, Turkey, in 2010 (31). However, in the end all efforts were in vain. The last bird disappeared in 2013 and with it also the last historic migration tradition (32). This event also marks the general extinction of the Northern Bald Ibis in its characteristic lifestyle as a migratory species. There is no longer any evidence that a migrating population still exists anywhere in the former distribution area.

What remains after vanishing of a species are cultural traces and these can be found throughout the entire historic area of this extraordinary bird. But the richest and most exciting traces are from ancient Egypt.

### The Akh

In ancient Egypt, the image of the Northern Bald Ibis was inseparably liked with the notion of *akh*, often translated as blessed dead or effective spirit (33–38), pointed toward many different meanings, such as the efficient blessed dead or living people who acted effectively for or on behalf of their masters (39). The *akh* belonged to cardinal terms of ancient Egyptian religion – almost as important as terms saint or angel in Christianity! – and hence it can be often found in Egyptian religious texts, as well as in other textual and iconographic sources.

Its basic meaning related to effectiveness and reciprocal relationship that crossed the borderlines between different cosmic spheres (38). The Egyptians considered their blessed, efficient and influential dead (the *akhu*) ‘living’ or ‘transfigured’, but a deceased human being had to be admitted and elevated into this new state. The dead became *akhu* only after mummification and proper burial rites were performed on them and after they had passed through obstacles of death and the trials of the underworld. The positive status of the mighty and transfigured *akhu* was mirrored by a negative concept of the *mutu* who represented those who remained dead, i.e. the damned.

A very important role in the process of becoming an *akh* was reserved for the horizon (*akhet*). Although it mainly represented the junction of cosmic realms (the earth, the sky and the netherworld), the horizon was a region in itself (40), and it was believed to be the place of sunrise, resurrection and a region where divine beings dwelled and where they interacted with the world of the living (33,41–44).

When a dead person’s journey to the afterlife had successfully finished and he/she was justified and transfigured into an *akh*, the person thus became a mighty and mysterious entity, which participated on the divine sphere of existence and yet still had some influence upon the world of the living. The *akhu* guarded their tombs where they promised to punish intruders on the one hand, and be inclinable to help those who presented them with offerings on the other (45), and acted as mediators who could intercede on behalf of the living with the gods or other *akhu* (37). Although the *akhu* had reached the afterlife existence, they still needed the living, since it was the latter who performed rituals, carried out the embalming and funerary requirements, and provided their dead ancestors with offerings (33). The *akhu* and the living represented co-dependent communities, and their mutual relationships and cooperation formed one of the pillars of ancient Egyptian religion (33).

### Appearance of the Northern Bald Ibis

The Northern Bald Ibis is an exotic appearance with a long curved bill, a naked reddish face, framed by an imposing crest of black lancet feathers, and an all-black plumage that gleams metallic in the sunlight (Fig 1). The Latin name reflects the characteristic appearance: *Geronticus eremita*, the old hermit. Systematically, the Northern Bals Ibis belongs to the order Pelecaniformes and the family Threskiornithidae. The genus *Geronticus* includes only further species, the Southern Bald Ibis (*G. calvus*) native to South Africa.

The sexes do not differ significantly. The males are slightly heavier than the females (mean weight m: 1390g, f:1257g; (10), with a longer and stronger beak. But the distributions of characteristics overlap, and a reliable determination of the sexes is only possible with genetic methods. The juveniles, on the other hand, are clearly distinguishable by their grey-feathered head.

Most historic Northern Bald Ibis drawing represent adult birds with the bare head and the feather crest (6,13,46). But there are some noticeable exceptions. A drawing by Conrad Gessner (11) shows a juvenile bird with a small crest and completely feathered head. Even more clearly is an altarpiece from the fifteenth century from around Munich, Germany, which represents the Mount of Olives scene, with Jesus and the disciples in prayerful attitudes. A detailed juvenile Northern Bald Ibis is shown at the edge of this picture, even with a worm in its bill as characteristic food. Such detailed representations indicate that the artists knew the birds themselves, what is an important indication for the presence of the species in these times.

The Bald Ibis in the altar scene is interpreted as a representation of death and the hereafter (4,13). Christian cultures in Europe regarded black birds in general rather as bad omen or nuisance (16). But the rather negative mythological image probably did less harm to the species than its reputation as a tasty food bird. Gesner (11) reported that the Northern Bald Ibis was praised as food and considered a treat because of the lovely flesh and soft bones. They were caught, shot and pre-fledged juveniles were taken from the nest (21).

The importance for the dining tables of the nobility and the clergy even led to protective measures as the populations declined. In Salzburg, for example, archbishop decrees of 1504 and 1584 criminalized shooting of Northern Bald Ibises in the wall of the *Mönchsberg* above the city of Salzburg. In the same century, Emperor Maximilian I in Graz provided artificial nesting aids in the rock walls. In the same period, an order was issued in the city of Graz in Austria, where also a colony occurred, that Northern Bald Ibises should not be shot, but rather cherished, controlled and guarded (24,47). Even these measures could not prevent extinction, but at least the last evidence for Bald Ibis occurrence in Europe comes from Graz in 1621 (21).

In the Muslim world the Bald Ibis had a better and more significant mythological image than in the Christian world and they were less pursued as edible birds. In Muslim tradition, the birds were worshiped because it was a Bald Ibis who led Noah and his family to the fertile lowlands on the Euphrates after landing on Mount Ararat (7). In Bireçik they also honoured these birds as leaders of the *hajj*, because they flew southward in fall in large numbers, just like pilgrims to Mecca, and they returned in spring after a period common for pilgrims (4,24). For that reason, the people of Bireçik used to celebrate the return of the birds in February with a traditional *Kelaynak* festival (*kelaynak* is the Turkish name for the species).

### Akh in material, textual and pictorial evidence

Ancient Egyptians used a pictorial representation of the Northern Bald Ibis for the hieroglyphic sign ‘*akh*’ (Fig 2). The sign, like its living model, is easily recognizable by the shape of its body, posture, shorter legs, long curved bill and a typical crest covering the back of the head. Although there are many aspects of this bird’s nature that must have had impact on the mind of the Egyptians, the main factor in holding the bird in particular esteem and relating it to concept of the *akh* was probably the bird’s habitat (see below).

**Fig 2.**
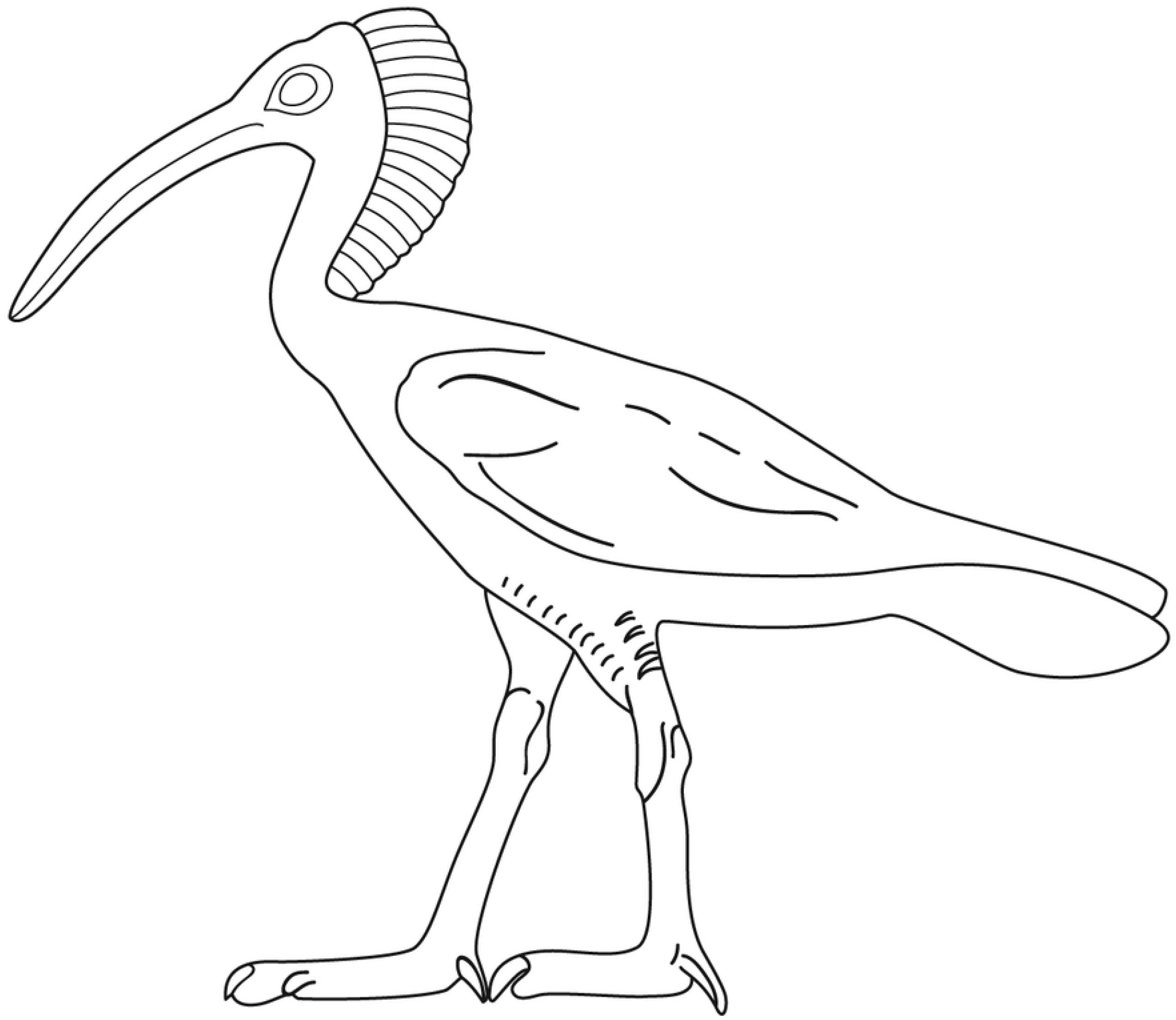
The hieroglyphic *akh*-sign from the tomb of Akhethotep. Drawing by Lucie Vařeková.

The only attested piece of material evidence for the Northern Bald Ibis in Egypt in the form of skeletal remains comes from Maadi region (48) located south of modern Cairo where the so-called Maadi culture had its settlements around 4000 – 3400 BC. This unique find represents both the earliest evidence for the presence of the bird in Egypt and its only confirmed preserved bodily remains. Pictorial representations of the Northern Bald Ibis have been recorded only from later periods of Egyptian history. The earliest Egyptian example of the bird’s depiction is attested on the so-called Ibis slate palette dated to the Naqada III Period (circa 3300 – 3000 BC), other early examples date to the Late Predynastic and Early Dynastic Periods (c. 3000 – 2700 BC).

From the Old Kingdom (c. 2700 – 2180 BC) onwards, a pictorial representation of the Northern Bald Ibis was used as a hieroglyphic sign for the word-root *akh* linked with the notion of effective power and the blessed dead. Some of the Old Kingdom depictions of the bird (e.g. from the tombs of Hetepherakhti, Akhethotep, Ptahhotep II or Ankhmahor) show very high accuracy revealing precise observations of ancient scribes and artists. On the other hand, depictions of this ibis attested in later tombs, e.g. the one of Hesuwer and Khnumhotep II dated to the Middle Kingdom (c. 2130 – 1770) are not as detailed as earlier examples. In Khnumhotep’s case, the Northern Bald Ibis is represented in a surprisingly incorrect manner (Fig 3), although other bird depictions attested in the tomb show unique accuracy and detail (8,49). The Northern Bald Ibis also appears on several Old Kingdom diadems found only in funerary context. These objects are equipped with additional discs composed of two opposed papyrus umbels with a Northern Bald Ibis on each of the blossoms. In some cases, an *ankh* (the sign of life) appears between the birds (50,51). The funerary context suggests that the diadems probably meant to ensure the reaching of afterlife and the ibis (as the *akh* sign/symbol) points towards the idea of transfiguration and resurrection (39).

**Fig 3.**
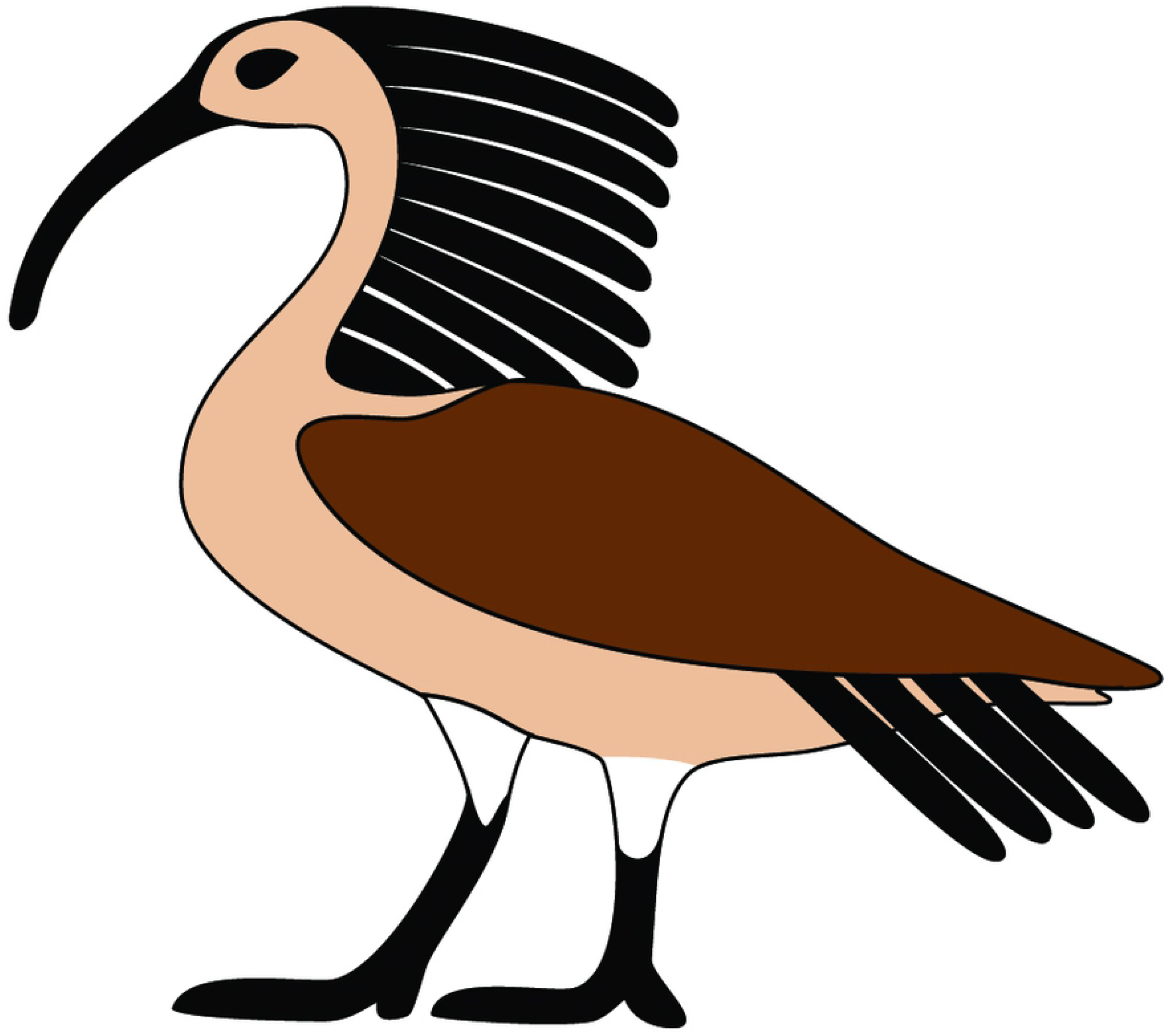
The hieroglyphic *akh*-sign from the tomb of Khnumhotep. Drawing by Lucie Vareková.

From the time of the New Kingdom (c. 1550 – 1070 BC) and the following periods of Egyptian history artistic representations of the Northern Bald Ibis are almost completely missing with two exceptions. The first is represented by the sign of *akh* that has kept its form (a rather stylized depiction of the Northern Bald Ibis) until the very end of ancient Egyptian history and the demise of hieroglyphic script. The latter deals with a mysterious ritual, called the *Vogellauf* in Egyptology. The ritual is attested among royal cultic scenes depicted mainly on temple walls from mid-New Kingdom until the Roman period (3,52,53) and which was most probably associated with another two still partly enigmatic rituals, called the *Rudderlauf* and the *Vasenlauf* (52). The representations of the ritual show the king running towards a deity with a Northern Bald Ibis in his one hand and three rods or sceptres of life, stability, and power in the other (Fig 4). The scene does not refer pictorially or textually to the ibis as to an offering or a sacrifice and the bird’s representation leaves us to conclude that it was neither dead nor alive. Hence, the suggested interpretation is that the image of the Northern Bald Ibis does not refer to the bird in its self but rather represents again the *akh*-sign in a symbolic reference to the king’s effective *akh*-power and mutual *akh*-relationship between the king and the deity (3). The loss of accuracy and the lack of evidence was very probably linked with the disappearance of the species from Egypt during the final phase of the third millennium BC (see below).

**Fig 4.**
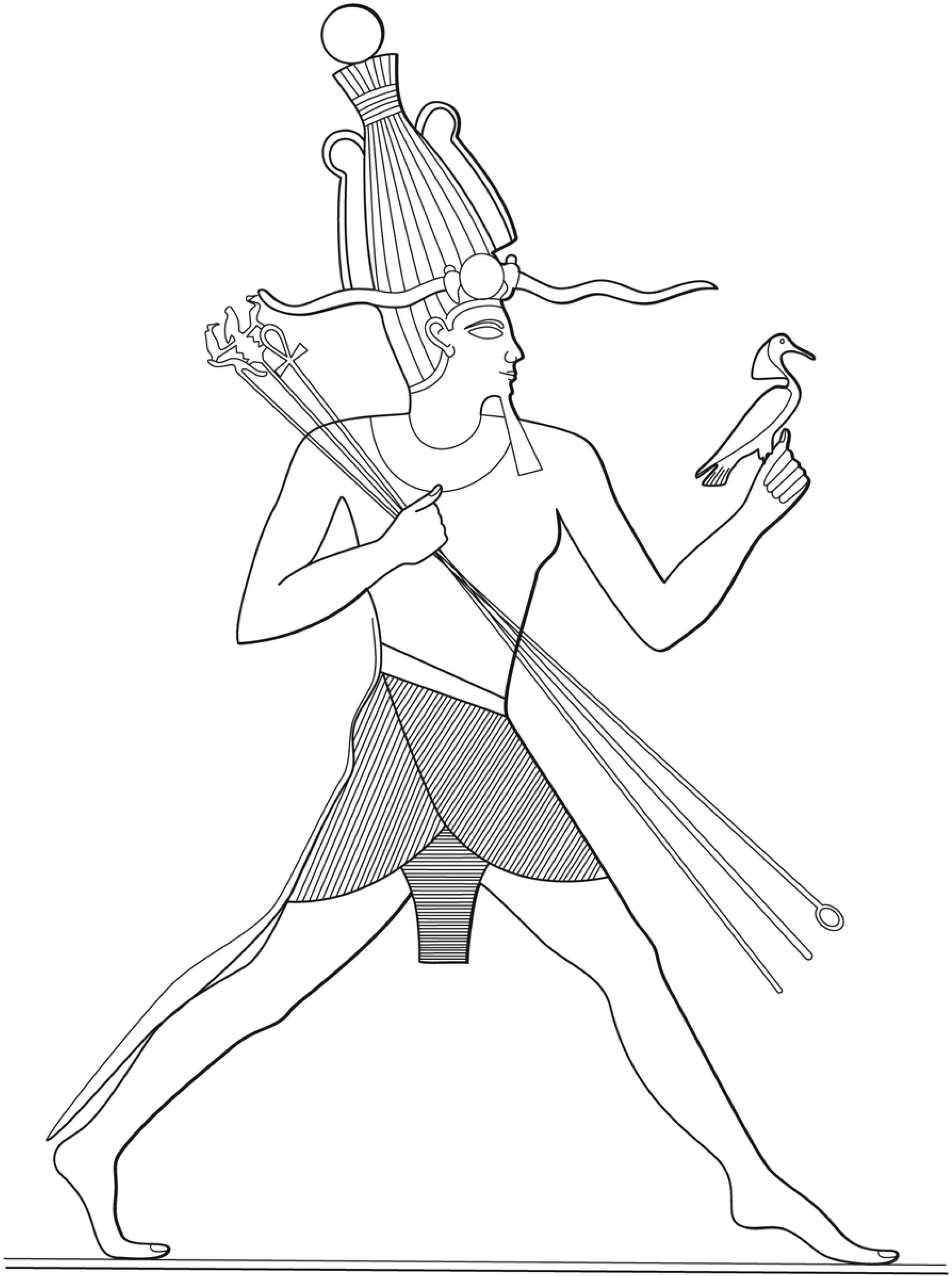
Egyptian king during the so-called *Vogellauf* ritual. Drawing by Lucie Vařeková.

### Northern Bald Ibis habitats

A remarkable characteristic of the Northern Bald Ibis is the long and fragile curved bill. It is poorly suited to hunt for mobile prey but perfectly shaped to dig for invertebrates deeply into the soil. Thus, the Northern Bald Ibis is mainly a tactile hunter. Favourable habitats are open landscapes with low vegetation and a high abundancy of the soil fauna. Under favourable conditions, the diet of the specie predominantly consists of worms and larva (54,55). But Bald Ibises show a high flexibility in their feeding habits. For example, in the Syrian desert birds were found to feed mainly on tadpoles which they pick up from the beach out of man-made reservoirs (56), while the same individuals at the wintering site predominantly dig for worms and larvae at freshly cut hayfields (57). For the remaining population in Morocco, lizards were found to represent an essential part of the diet (58).

The flexibility in foraging is also reflected in the diversity of foraging habitats. Northern Bald Ibises used to feed in the Syrian desert (59), in the Moroccan steppe (58,60), on meadows and pastures of the northern foothills of the Alps (54,55) or the Ethiopian highlands (57) and even on Turkish mint fields (61). Feeding habitats have in common that they are rather open landscapes with at least a loos vegetation coverage. The birds show a clear preference for low vegetation, usually not higher than 10 cm. This can be a natural characteristic of the vegetation, especially in semi-arid areas, but in most region the birds benefit from grazing or mowing (55,60). Moreover, the Northern Bald Ibis is generally not a bird of wetlands, as many other ibis species, but it needs an available freshwater source for drinking and bathing. This became evident in Morocco, where the provision of freshwater near the breeding colony significantly enhanced breeding productivity (62).

A noticeable peculiarity in connection with the feeding ecology is the frequent proximity of the breeding habitats to human settlements. From historical reports this is evident for the former European population (11–13,16). It is also known for most former breeding sites in Moroccan and Algerian Atlas (6,60), in Turkey (25,27) and for the former wintering site in Ethiopia (57). It is assumed that the presence of the species in various regions was dependent on human beings which cleared or drained the land and kept it open through farming or grazing (12,13,16,60). This can be regarded as a kind of mutualism, because the birds have benefited from the open sites and have eaten larvae of pest insects for their part. But as with many other species, proximity to humans ultimately also carries a high risk for the species depending on the respective culture and period.

Also, the breeding sites were often close to human settlements. But this was probably more of a coincidence than a mutualistic relationship. The Northern Bald Ibis as a colonial breeder needs cliffs which are structured with niches or ledges (Fig 5). Many such cliffs consisted of limestone or conglomerate, located on the sea, along large rivers or on lake shores. And these were often also preferred areas for establishing human settlements. Examples are the former Turkish colony in Bireçik at the river Euphrates (25, Fig 6) or former European colony sites like Salzburg on the river Salzach (Fig 7), Passau on the river Danube or Uerberlingen at Lake Constance (12). This coincidence also contributed to the extinction, indirectly through disturbance and destruction of the breeding sites, but also through hunting and collection of nestlings out of the nests.

**Fig 5.**
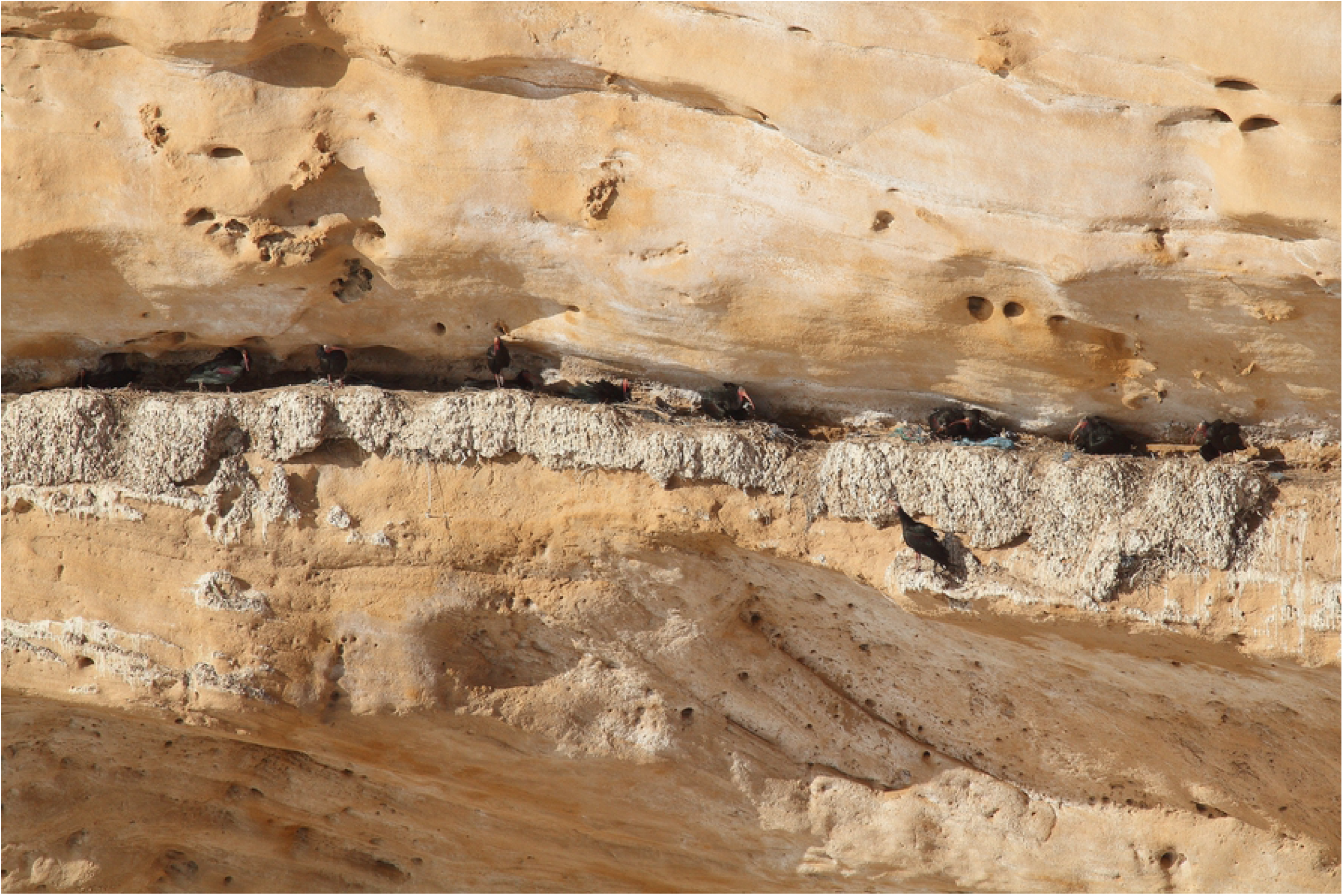
Breeding cliff in Agadir, Morocco. Picture D. Tome.

**Fig 6.**
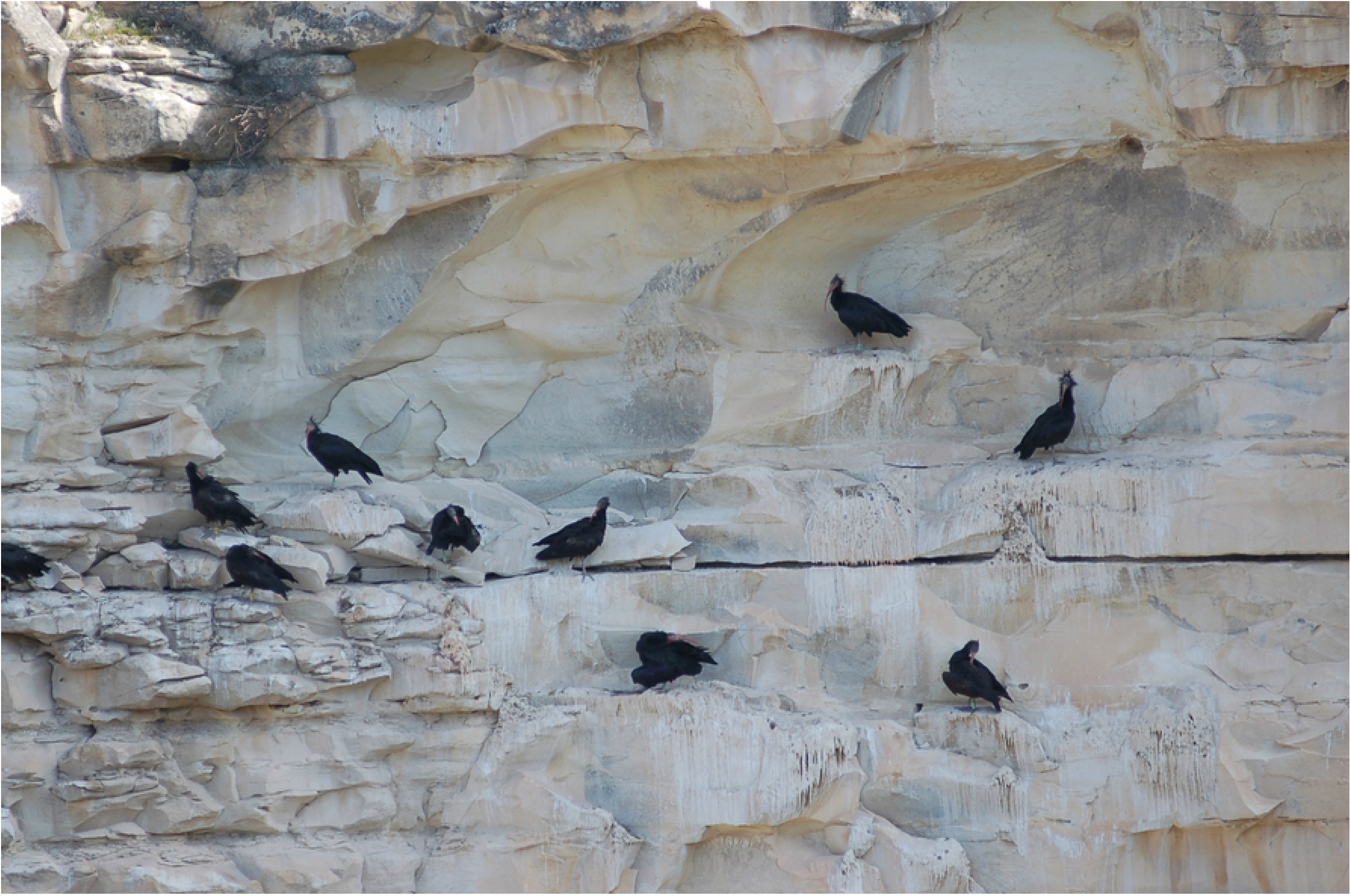
Breeding cliff in Birecik, Turkey. Picture J Fritz.

**Fig 7.**
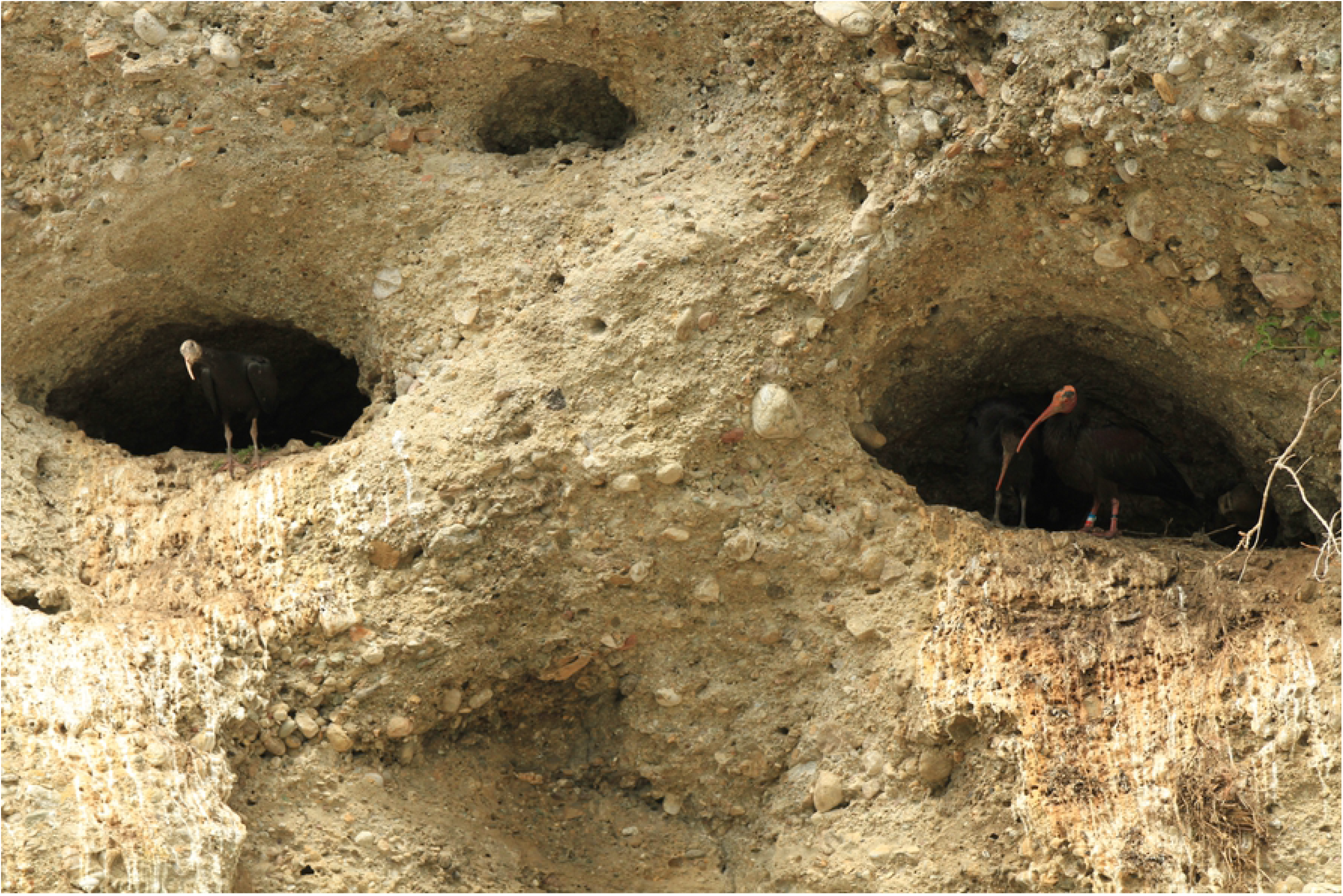
Breeding cliff in Kuchl, Austria. Picture J Fritz.

### Akh and the eastern horizon

From the point of view of ancient Egyptian cosmology and religion, the most important place linked to the idea of the *akh* and related notions was the horizon. Not only did the Egyptian term for the horizon (*akhet*) share the word-root *akh*, but it also cohered with the ideas connected with the afterlife and the divine sphere. The horizon represented the junction three main cosmic realms, the earth, the sky and the underworld, and as a place of sunrise it was also considered to be the symbolic place of birth, renewal and resurrection. The blessed deceased (the *akhu*) who overcame the obstacles of death and successfully ended their journey through the underworld at the eastern horizon of heaven.

The eastern horizon was, thus, closely linked with the *akhu* who were believed dwell at and come from the *akhet* (i.e. the eastern horizon). In fact, ancient Egyptian sources witness that the *akhu* are ‘born’ or ‘created’ in the *akhet* and their often refer to the blesses deceased as to those “who dwell at the horizon” (39,43,44). The Egyptian hieroglyphic sign of the (eastern) horizon had a shape of rocky peak or rather a cliff massive with two peaks. Throughout the Nile valley, the eastern horizon is in fact created by limestone cliff massif with occasional peaks. And knowing how precise the Egyptians were in observing nature, it is not by accident that the transfigured blessed deceased were denoted *akhu* and that the hieroglyphic signed used for them was a picture of the Northern Bald Ibis: the birds and the dead occupied the same habitat.

### Characteristic Northern Bald Ibis behaviours

The Northern Bald Ibis is a year-round social species with hierarchically structured colonies consisting of up to several thousand individuals. The species is monogamous, where some couples form long-lasting bonds while others stay together only for a breeding-season. Both partners breed and raise the chicks, as it is characteristic for many monomorphic species (63). Fledging rate varies considerably between the populations, ranging from 1.0 to 2.2 chicks per nest dependent on site and condition (10). According to data from the European release project, the family groups already dissolve in the breeding area. The juveniles join together in groups which follow experienced conspecifics to the wintering site (64).

The species has a rather moderate repertoire of calls. Most common is the ‘*croop’* call, which is used in an affiliative context - the characteristic greeting behaviour with the rhythmic vertical movement of the beak - as well as during agonistic encounters (65). The ‘*croop*’ was found to have highly variable temporal and structural parameters, which may indicate the expression of affective states and even encode individual differences that allow individual recognition (66). This results in parallel with a morphological characteristic of the species, the conspicuous bare head. In adult birds, the pattern of the dark areas allows individual recognition, even for humans (67). Moreover, in males the bare throat area varies in size between individuals and has a seemingly hormonally controlled variation in the intensity of the red colouration (68). According to that it is assumed that the bare head of the Northern Bald Ibis evolved mainly in the context of social interaction and mate choice. However, there is also some evidence that in species like the Northern Bald Ibis inhabiting hot environments bare skin has evolved to dissipate heat (69).

A noticeable and characteristic behaviour of the Northern Bald Ibis, as of other Ibises (*Threskiornithidae*), is the sunning resp. sun-bathing behaviour, where the bird remaining in a stiff upright position with wings outstretched (Fig 8). This behaviour, also known as sun worship, has contributed to the veneration of this species in various cultures (7). However, the actual function of this behaviour is still unclear (70). Hypotheses are thermoregulation, killing of ecto-parasites by heating the feathers and the body surface or vitamin D-synthesis by exposition of bare skin on the underside of the wing to the sun. In any case, sunning behaviour has a clear social component, when a bird starts with it, it usually stimulates other conspecifics, which makes this behaviour even more conspicuous.

**Fig 8.**
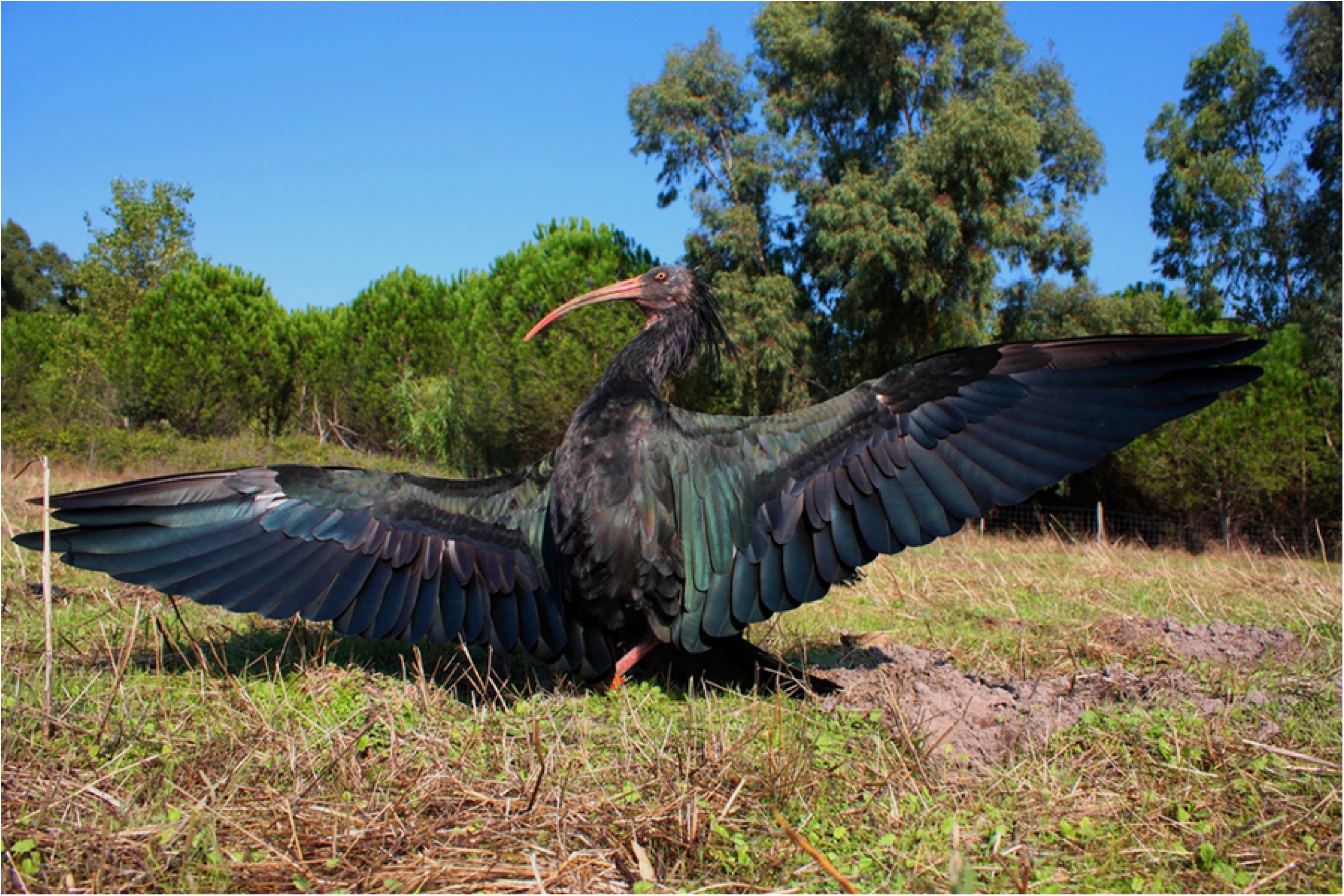
Sunning behavour. Picture M. Unsoeld.

### Social aspect of the *akh*

The *akhu* had a unique position within the ancient Egyptian concept of the natural world. The highest position among all cosmic beings was held by the gods and the lowest was reserved for human beings, but the boundaries between the human and divine worlds were occupied by semi-divine entities with super-natural status and power, mainly the blessed and the damned dead. If a person’s underworld journey has successfully finished with justification and acceptance into the afterlife, such person was transfigured into an *akh*, and thus became a mighty and influential entity with supernatural powers who still had some influence upon the world of the living. The *akhu* guarded their tombs, would on the one hand punish intruders, and on the other help those who made pleas to them. The *akhu* would help the living in cases when human abilities were insufficient. The dead, however, still needed the support of the living, who performed rituals, mummification and funerary rites, and, most importantly, the living provided their dead ancestors with offerings. The *akhu* and the living represented co-dependent communities. The mutual relationship of these two communities of cosmic beings formed one of the main pillars of ancient Egyptian religion (37,39).

It was this bilateral *akh*-efficiency of reciprocal actions what helped to cross the threshold of death and bridged the boundaries of this world and the afterlife. The *akhu* represented by the Northern Bald Ibis both in iconography and in the real world held the role of influential intermediators and guarantors of survival. For Egyptians of lower social strata, the *akhu* were even more important than the gods.

The society of the blessed dead had also its own hierarchy structured similarly to the world of living. At the very top of the *akhu*, there was only one person: the deceased king. According to the Pyramid Texts (e.g. § 833, 858, 869, 899, 903), the Egyptians called him ‘the head of the *akhu*’ or ‘the first of the *akhu*’ or even ‘the *akh* of the *akhu*’. Like in the real world, also the society of the *akhu* had a pyramidal structure. Below the sole position of the king there were ‘successful and well-equipped *akhu* (of the sun god)’ who often recruited from the strata of elite officials. For the non-elite Egyptians, however, deceased family members and local patrons represented the most important allies in the afterlife. The strictly hierarchical structure of the *akh* society with only one entity at the very top may have not only reflect the structure of human society but may have also stemmed from the well-observed V-formation used by the birds during migration (64).

### Northern Bald Ibis migration

The Northern Bald Ibis was a migratory species all over the historic range with known wintering sites along the African west coast down to Mauritania and Mali and along the African east coast down to Ethiopia and Eritrea (29,58,71). For the former European population, it is know that they left in autumn and returned in spring, but any evidence for the migration pathway or the historic wintering site is lacking (12,13). The migratory European release population migrate to a wintering site in the southern Tuscany. However, this sites was selected for the release not on the basis of historic records, but rather due to the current and sustainable suitability of the habitats (71).

Under appropriate ecological and climatic conditions colonies of various migratory species are known to shorten their migration route or change to a resident lifestyle. This happens especially along coastlines with year-round moderate climatic conditions (72). This also applied for the Northern Bald Ibis. Several resident colonies are known along the Atlantic coast in Morocco, but such colonies probably also existed along the Red Sea. Meanwhile, all migrating colonies have been eradicated, while two residential colonies on the Atlantic coast still exist (58,60).

Due to extinction of all migratory colonies most knowledge about the species-specific migration behaviour mainly comes from the migratory European release population. These birds are descendants of former migratory colonies in the Moroccan Atlas. Research on physiology, energetics and behaviour has shown that the Northern Bald Ibis is an enduring migratory species with a pronounced navigation ability. In Europe, the birds enter into a migratory state (Zugbereitschaft) beginning of August (73,74). At that time, they leave their breeding areas and usually return in the next season from the end of March (75). The Middle East colonies (mainly Bireçik and Palmyra) hat their rhythm shifted by about one month with departure in early July and return from the end of February (10,29).

Northern Bald Ibises are persistent flyers with a daily migration flight stages of up to 350 km (30,31,75) and an average flight speed of about 45 km/h (76). They use different flight techniques to save energy. The most noticeable one is the V-shaped or echelon formation, where energy savings can be achieved by using the aerodynamic up-wash produced by the preceding bird (77). As the leading bird in a formation cannot profit from this up-wash, a social dilemma arises around the question of who is going to fly in front. The Northern Bald Ibises solve this dilemma by directly taking turns in leading the formation (78). This is assumed to be one of the rare examples of real cooperation in the animal kingdom (79).

### *Akh* migration

Although have almost no physical evidence on the presence of the Northern Bald Ibis in ancient Egypt and on the nature of its stay there, we may strongly presume that the bird was only a migratory visitor to the country on the Nile. This is based upon the already discussed nature of the bird, character of its Egyptian habitat (see above) and – strangely enough for some – upon ancient religious texts. The Egyptian connected the deceased with migratory birds whom they saw as messengers from the realm of the dead. In a famous line attested in royal tombs of the New Kingdom, migratory birds are described as beings with avian bodies and human heads coming to Egypt from the North, from the region of utmost darkness (80). However, no direct evidence of the possible migration of the Northern Bald Ibis to, from or via Egypt has not yet been attested. In this respect, there are questions that need to be answered.

Does the bird’s Egyptian habitat (limestone cliffs) provide us with any clue to the nature of its presence in the country and its possible migration? Does any of the three seasons of the Egyptian year (the season of high inundation, the season of growth and the season of low water) fit more to the needs of the bird? Was the appearance of the *Akh* star/constellation on the night sky a particular period of the year somehow connected with the arrival or departure of Northern Bald Ibises to/from the country? The problem still needs to be studied, as its solution may shed a new light on many aspects of ancient Egyptian view of the natural world as a sacred space. Close multidisciplinary cooperation on the topic would give us the best change to solve the problem.

### Endangerment and disappearance of the Northern Bald Ibis

In 2018, after 24 years, the species was down-listed on the IUCN Red List from *critically endangered* to *endangered*. This was mainly justified by successful conservation measures to secure the last wild population in Morocco. However, this decision was controversial. Although this population has developed well in recent years and signs of expansion of the breeding area have even been observed (10,81), it remains so far the world’s only wild population, spatially restricted to a small area on the Atlantic coast in Morocco. The significant decline of the population in 1996 as a result of an epidemic (82) and the dependence of the population on management like the provision of supplementary fresh water (62) indicate that this one population cannot ensure permanent survival of the species.

Accordingly, the major purpose of the International *Single Species Action Plan for the Conservation of the Northern Bald Ibis*, published in 2015 and foreseen for a ten-year period (83), is to increase the population size and the breeding range of the species. This should mainly be achieved by improved management of the existing Moroccan colonies and establishment of new colonies, with a particular focus on former breeding area outside Europe, where colonies recently disappeared.

Though, by the halfway point of the action plan no concrete translocation projects had been initiated in these regions. This has mainly economic, logistical or political reasons but is also related to the fact that former causes of extinction are still present in these areas, in particular uncontrolled hunting, electrocution, poisoning and the ongoing destruction of habitats (29,32,84).

Meanwhile, two successful reintroduction projects are being implemented in Europe. *Proyecto Eremita* in Andalusia, Spain, is on the way to establish a residential population which consisted of around 140 individuals end of 2019 colony (10). The birds breed at three side in an area of 23 km along the Atlantic coast in Andalusia. Every year around 40 juveniles from various European zoo breeding colonies are supplemented to build up a self-sustaining population (M Quevedo, pers.com.).

At the same time *Waldrappteam* establishes a migratory colony in Central Europe (4,71) with 142 birds end of 2021, divided into three breeding colonies north of the Alps and one further colony in Carinthia, Austria. These birds migrate along two distinct migration corridors between the breeding sites and the wintering site in southern Tuscany. The project became famous because of the human-led migration, where human-raised and trained juveniles follow their foster parents in two microlight planes from the breeding sites to the wintering area, where they will then be released (Fig 9). A population viability analysis indicates that the population needs a minimum size of about 320 individuals for self-sustainability (85). This threshold should be exceeded in the mid-2020s. Modelled scenarios also indicate that this European population is relatively stable against stochastic events that can be caused, for example, by climate change (85).

**Fig 9.**
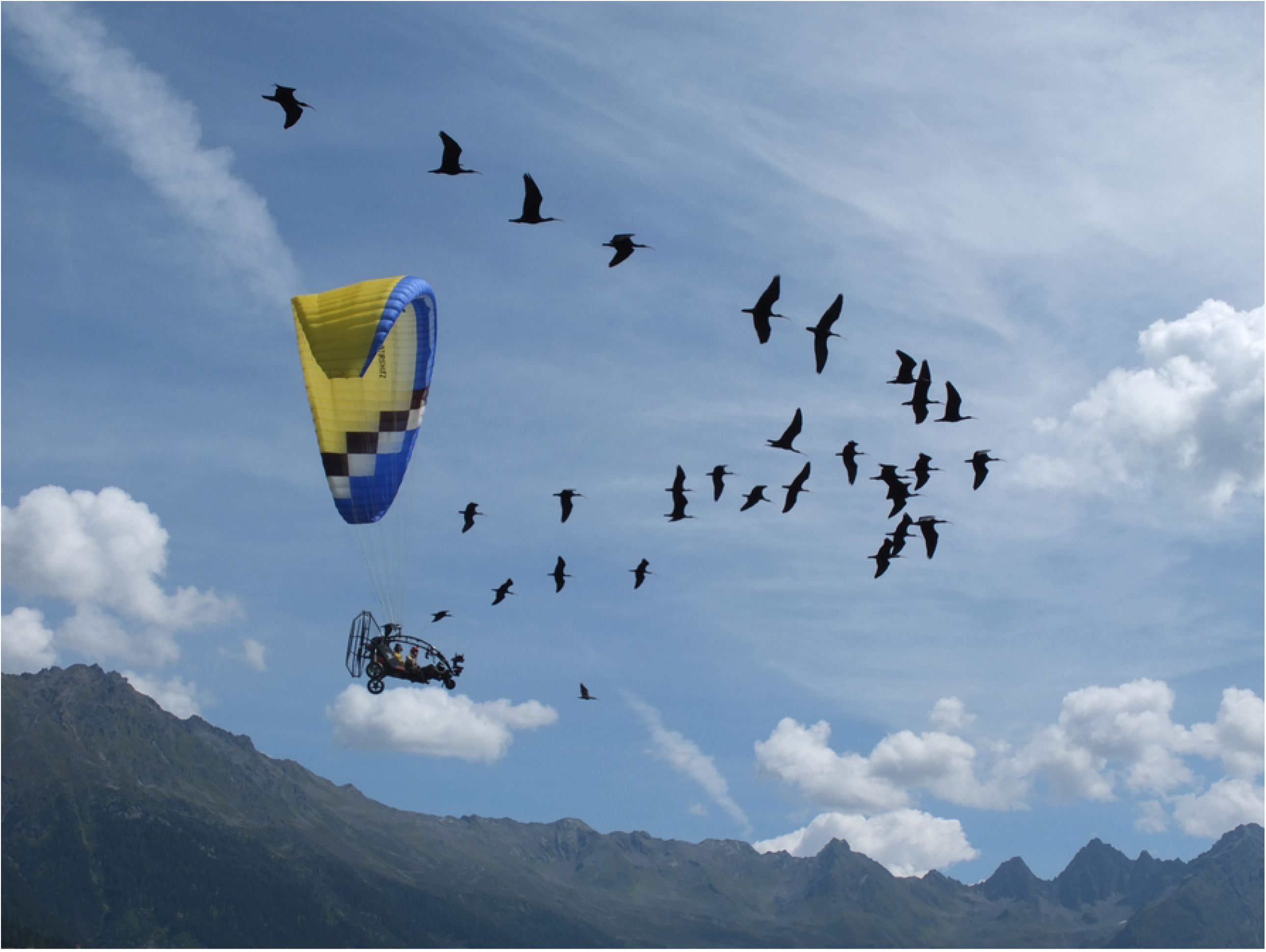
Human led migration flight across the Austrian Alps. Picture C. Esterer.

### Akh disappearance

As has already been discussed above, we lack physical evidence on the presence of the Northern Bald Ibis in Egypt. Thus, unfortunately, one can only use pictorial and textual evidence in researching the nature of the presence of the bird in the country. In this respect, a particularly noteworthy fact can be revealed by analysis of the Northern Bald Ibis iconographic features. Pictorial evidence on the bird is much more accurate, precise, and elaborate in the early periods of Egyptian history, until the final phase of the third millennium BC (8). In later times, the representations of this ibis become very schematized, sometimes they even do not correspond to their natural model very much. They cannot be viewed as convincing evidence for the presence of the Northern Bald Ibis in Egypt. Moreover, there is no material, pictorial, or textual evidence for keeping, breeding, hunting, killing, mummifying, or sacrificing the Northern Bald Ibis in ancient Egypt from any period of its history. We have no evidence for hunting or sacrificing the bird in Egypt, nor was it kept in temples and mummified at death (48,86). The last mentioned fact stands in striking contrast to the Sacred Ibis and the Glossy Ibis (*Plegadis falcinellus*) that are known to have been kept and mummified (48,87); there are many thousand mummified examples of the Sacred Ibis (*Threskiornis aethiopicus*) (86).

Judging from iconographic evidence, at the latter phase of the Old Kingdom (towards the end of the third millennium BC), the Northern Bald Ibis has begun to disappear from Egypt, or rather it has altered its migration routes, avoiding Egypt at all or making only a few stops there, or that the present migration route of the Northern Bald Ibis from Ethiopia to Syria that avoids Egypt (30) already originated at the beginning of the second millennium BC (8). By why this change occurred?

Here again we face a problem that can only be solved by a close multidisciplinary nature-human science cooperation. At the present stage of research, we are able to ascertain that the time of the disappearance of the Northern Bald Ibis from ancient Egypt follows a climate change period: a period of a swift desiccation of the country and expansion of arid areas that occurred in the first half of the third century when many other animal and avian species left the country (88–90). But unlike the Elephant, the Giraffe or the Saddle-Billed Stork who disappeared from Egypt during the time of the climate change and a gradual desiccation, the Northern Bald Ibis is leaving only 500 years later.

Data gained from a case-study made on the Saddle-billed Stork (*Ephippiorhynchus senegalensis*) can well serve as a comparative material for studies on the disappearance of the Northern Bald Ibis. Similarly to the Northern Bald Ibis, the Saddle-billed Stork was also closely connected with a very important religious concept, as its hieroglyphic depictions served to denote ‘divine power’ or a ‘manifestation of divine power on earth’ (the *ba*). The study (8,91) proved that after the species disappeared from Egypt after a climate change early in the third millennium BC, first the hieroglyphic sign lost its accuracy and slowly stopped to resemble the stork at all, then the sign and Egyptian term *ba* was given a new meaning (the soul) and, finally, a completely new hieroglyphic sign (a human-headed falcon) for the *ba* was invented. The study on the stork clearly shows that the Early Dynastic climate change had a direct impact not only upon nature, human life and society, but also upon such seemingly unrelated phenomena as script, religion and philosophy (91).

The most important question of the present article still stands. Why the bird left the country when there were no dangers like hunting or pesticides that have endangered the species in modern times, nor there was no big harm made to it by the climate change itself? Although the research on the topic is at the very beginning, we can say that the most important factor most probably was human activity in and around the bird’s feeding and breeding areas.

The time of the disappearance of the Northern Bald Ibis from Egypt was the period of (I) higher human activities in the fields following the need for new irrigation projects after a climate change, (II) higher building activity (the species disappeared during the so-called age of the pyramid builders), which lead to a higher human activity in the bird’s breeding region at the limestone cliffs that were used as quarries) and (III) turbulent human activities linked to social disorder that occurred after the collapse of the Egyptian state at the end of the Old Kingdom, probably originally also linked with earlier climate change.

## Discussion

Both in Egyptology and in Ornithology, the Northern Bald Ibis represents an iconic bird species. Coincidently, it is connected with the notion of death and disappearance in both scientific fields. In ancient Egypt, the Northern Bald Ibis was linked with the concept of the blessed dead (mighty spirits called the *akhu*) who once disappeared from the land of the living but only to return there after a successful journey through the underworld (39). In ornithological and conservation science, the bird is best known for its endangerment, which almost lead to its total extinction, and for the present-day attempts to save the species and to reintroduce it back to nature (71).

This study is an outcome of a cross-disciplinary cooperation of the two authors and their respective scholarly fields. The research was undertaken simultaneously in the two scientific fields, each using methodology of the appropriate scholarly domain, with frequent consultations and mutual sharing of data.

The original aim was to broaden our modern understanding of historic notions, concepts and approaches using up-to-date data on this species, which the ancient Egyptians used to describe the religious concept in focus. Such Ornithology-to-Egyptology transfer of knowledge enriched Egyptology with information about the species’ shape, colouring, habitat, social habits, migration periods and migration routes and, thus, proved to be very efficient. And with this new data, only 200 years after Egyptology as a scientific field was born, we were able to clarify our modern view of one of the key Egyptian religious concepts (the *akh*) who helped to shape human thought in Egypt for more than three millennia. Only when one closely examines the model bird and learns about the habitat, migration or social behaviour of the Northern Bald Ibis, one is able to understand the ideas hidden in the background of the concept of the *akh* (8).

However, an in-depth Egyptological study on the concept of the *akh* and on its hieroglyphic sign proved to be valuable for Ornithologists as well. First, after gaining information from ancient Egyptian texts (some of them more than 4500 years old), Egyptologists were able to discern the V-shaped flight formation of the Northern Bald Ibis as an important piece of evidence for the migration behaviour of the species in this region in ancient times. The most important result of the Egyptological part of the research on the Northern Bald Ibis was represented by the gaining of significant data to show that the bird had disappeared from Egypt during late third millennium BC after a climate change (8). With this research step the Egyptology-to-Ornithology transfer of data, information and knowledge began or became stronger. Not only one can follow the bird’s presence, habits and habitats almost 5.000 years back to history of the world, but the evidence is formed by textual and pictorial materials, not only by bones and other bodily remains.

Ancient Egyptian sources thus present us with valuable information both on the presence of the Northern Bald Ibis in the Egyptian Nile Valley and Delta and on its early disappearance. But what is of utmost importance is the fact that these ancient sources witness an indirect link of the bird’s disappearance from Egypt to the climate change.

The bird disappeared from Egypt more than 500 years after the climate change, as the reasons for the disappearance were rather connected with changes in human behaviour resulting from the climate change effects. The time of when the Northern Bald Ibis began to leave Egypt was the period of higher human activities in the bird’s feeding area (following the need for new irrigation projects after a climate change), increased quarrying and building activities in the bird’s breeding areas (the age of the pyramid builders), and last but not least, turbulent human activities stemming from social disorder after the collapse of the Egyptian state at the end of the Old Kingdom (this socio-political phenomenon was also originally connected with the earlier climate change).

Climate change is also an increasing threat to the last remaining wild population in Morocco. Morocco is among the countries which are expected to have the strongest effects in terms of temperature-rise, decreased of precipitation and weather extremes and the coastal region is assumed to be particularly affected (92). This will put increasing pressure on the Northern Bald Ibis population there. Under these changing conditions the residential lifestyle of this coastal population can become detrimental because residential populations lack ecological flexibility related to migration behaviour and there is hardly any evidence that residential populations are able to change back to a migratory lifestyle (72,79,93–95).

Noteworthy in this context, the *Single Species Action Plan for the Conservation of the Northern Bald Ibis* (83) don’t considers climate change, the term doesn’t even show up once in the whole text (83). This although the ecosystems of the majority of these areas are assumed to be disproportionally affected by climate change effects (96). History teaches us that Northern Bald Ibis populations can be significantly affected by the consequences of climate change. Therefore, regarding the purpose of the Action Plan to re-colonize former habitats, feasibility study should include modelling to examine whether newly established colonies can be sustainable with respect to climate change effects and related stochastic events. Such a feasibility study should also differentiate between scenarios for migratory and sedentary colonies as we have knowledge of the differences in ecological flexibility due to these different lifestyles (85).

Future interdisciplinary research shall focus the forms of human activities connected to the bird’s decline in more detail. This should endow natural scientists and conservationists with data that could support future conservation and reintroduction attempts with the Northern Bald Ibis and other endangered species, which is the highest goal of the two scholarly fields. Then the Northern Bald Ibis and the ancient Egyptian *akh* would again share the same meaning: a living entity that was gloriously resurrected after being dead.

## Conclusion

Tracing the history of the Northern Bald Ibis for several millennia provides us with some significant information about the coexistence of these bird species and humans in different epochs and regions. Coexistence in the form of a mutualistic relationship has worked well for long periods of time. But at some point, the situation changed to the disadvantage of the birds. Interestingly, significant changes that ultimately led to the extinction of Northern Bald Ibis populations were repeatedly accompanied by marked changes in the climate. This applies to the extinction of the population in ancient Egypt as well as of the European population in the Middle Ages and climate change also currently poses an increasing threat to the last remaining wild population in Morocco. The historical context indicates that we need to pay special attention to climate-change related effects in conserving and reintroducing this species. The interdisciplinary approach also illustrates that the conservation of the Northern Bald Ibis is also of significant cultural importance, not only because of historic meaning of this mythical species in different cultures but also to ensure that the Northern Bald Ibis can continue to enrich human culture in the future.

## Acknowledgments

We are grateful to Markus Unsoeld (Bavarian State Collection) and Katharina Huchler (Waldrappteam Conservation & Research) for discussions and helpful comments on the manuscript.

## References

1. Bennett NJ, Roth R, Klain SC, Chan KMA, Clark DA, Cullman G, et al. Mainstreaming the social sciences in conservation. Conserv Biol. 2017;31(1):56–66.

2. Campbell LM. Overcoming obstacles to interdisciplinary research. Conserv Biol. 2005;19(2):574–7.

3. Janák J. Running with images: Ritualized script in the Vogellauf, Rudderlauf and Vasenlauf. In: M. Bárta JJ, editor. Profane Landscapes, Sacred Spaces. Sheffield: Equinox; 2020. p. 89–96.

4. Fritz J, Unsoeld M, Voelkl B. Back into European Wildlife: The Reintroduction of the Northern Bald Ibis (Geronticus eremita). In: Kaufman A, Bashaw M, Maple T, editors. Scientific Foundations of Zoos and Aquariums: Their Role in Conservation and Research. Cambridge University Press; 2019. p. 339–66.

5. Fritz J, Unsoeld M. Artenschutz für einen histrosichen Schweizer Vogel: der Waldrapp im Aufwind. Wildbiologie Int. 2011;5(17).

6. Pegoraro K. Der Waldrapp. Wiesbaden: AULA-Verlag; 1996.

7. Hirsch U. Die Rettung der heiligen Vögel. Tierpark. 1976;9:4–11.

8. Janák J. Spotting the Akh. The Presence of the Northern Bald Ibis in Ancient Egypt and Its Early Decline. J Am Res Cent Egypt. 2011;46:17–32.

9. Bowden CGR, Aghnaj A, Smith KW, Ribi M. The status and recent breeding performance of the critically endangered Northern Bald Ibis Geronticus eremita population on the Atlantic coast of Morocco. Ibis (Lond 1859). 2003;145(3):419–31.

10. Boehm C, Bowden C, Seddon P, Hatipoglu T, Oubrou W, el Bekkay M, et al. Northern Bald Ibis: History, current status, and future perspectives. Oryx. 2020;(in press).

11. Gesner C. Historiae animalium liber III, qui est de avium natura; erste deutsche Übersetzung. Zurich: Christoffel Froschouer; 1557.

12. Schenker A. Das ehemalige Verbreitungsgebiet des Waldrapps Geronticus eremita in Europa. Der Ornithol Beobachter. 1977;74:13–30.

13. Unsoeld M, Fritz J. Der Waldrapp-ein Vogel zwischen Ausrottung und Wiederkehr. Wildbiologie 2 1-16. 2011;2:1–16.

14. Kumerloeve H. Waldrapp, Gerontocus eremita (LINNAEUS 1758) und Glattnackenrapp, Geronticus calvus (BODDAERT 1783): Zur Geschichte ihrer Erforschung und zur gegenwärtigen Bestandssituation. Ann Naturhistor Mus Wien. 1978;81:319–49.

15. Sánchez I. Evidence of the historic presence of the Northern Bald Ibis (Geronticus eremita) in Spain. 2006.

16. Perco F, Tout P. Notes on recent discoveries regarding the presence of the Northern Bald Ibis Geronticus eremita in the Upper Adriatic Region. Acrocephalus. 2001;22(106–107):81–7.

17. Boev Z. Presence of Bald Ibises (Geronticus Wagler 1832) (Threskionitidae (sic) - Aves) in the Late Pliocene of Bulgaria. Geol Balc. 1998;28(1-2):45–52.

18. Hölzinger J.Waldrapp (Geronticus eremita): Knochenfunde aus der spätrömischen Befestigung Sponeck am Kaiserstuhl. OrnJhBaden-Württemberg. 1988;57–67.

19. Mourer-Chauviré C, Philippe M, Guillard S, Meyssonnier M. Presence of the Northern Bald Ibis Geronticus eremita (L.) during the Holocene in the Ardèche valley, southern France. Ibis (Lond 1859). 2006;148(4):820–3.

20. Wirtz S, Boehm C, Fritz J, Kotrschal K, Veith M, Hochkirch A. Optimizing the genetic management of reintroduction projects: Genetic population structure of the captive Northern Bald Ibis population Sarah Wirtz. Conserv Genet. 2018;19(4):853–64.

21. Schenker A. Der Waldrapp - ein historisches Wildbret. Wildbiologie. 1981;4(7):1–12.

22. Kinzelbach RK. The distribution of the serin (Serinus serinus L., 1766) in the 16th century. J Ornithol. 2004;145(3):177–87.

23. Bauer KM, Blutz von Blotzheim UN. Handbuch der Vögel Mitteleuropas. Frankfurt am Main: Ak. Verlagsgesellschaft; 1966.

24. Peter H. Waldrappdämmerung am Euphrat. Heidelberg: Max Kasparek Verlag; 1990.

25. Hirsch U. Der Waldrapp Geronticus eremita, ein Beitrag zur Situation in seinem östlichen Verbreitungsgebiet. Vogelwelt. 1980;101:219–36.

26. Hatipoglu T. Conservation Project, Birecik, Turkey. Report of 4th IAGNBI Meeting Seekirchen (Ed C Boehm & C Bowden). 2016;40–6.

27. Yeniyurt C, Oppel S, Isfendiyaroglu S, Özkinaci G, Erkol IL, Bowden C. Influence of feeding ecology on breeding success of a semi-wild population of the critically endangered Northern Bald Ibis Geronticus eremita in southern Turkey. Bird Conserv Int. 2016;1–13.

28. Serra G, Abdallah MS, Assaed A, Abdallah A, al Qaim G, Fayad T, et al. Discovery of a relict breeding colony of northern bald ibis Geronticus eremita in Syria. Oryx. 2004;38(2):1–7.

29. Serra G, Lindsell JA, Peske L, Fritz J, Bowden CGR, Bruschini C, et al. Accounting for the low survival of the Critically Endangered northern bald ibis Geronticus eremita on a major migratory flyway. Oryx. 2014;49(2):312–20.

30. Lindsell JA, Serra G, Peške L, Abdullah MS, Al Qaim G, Kanani A, et al. Satellite tracking reveals the migration route and wintering area of the Middle East population of Critically Endangered northern bald ibis Geronticus eremita. Oryx. 2009;43(3):329–35.

31. Fritz J, Riedler B. Neue Hoffnung für das Überleben einer hoch bedrohtesten Zugvogelart im Mittleren Osten: Freisetzung von Jungvögeln bei den letzten migrierenden Waldrappen in Syrien. Vogelwarte. 2010;48:417–8.

32. Serra G. The Northern Bald Ibis is extinct in the Middle East - but we can’t blame it on IS. Ecologist [Internet]. 2015;(May). Available from: The Northern Bald Ibis is extinct in the Middle East - but we can’t blame it on

33. Friedman F. On the Meaning of Akh in Egyptian Mortuary Texts (doctoral thesis). Brown University; 1981.

34. Friedman F. A Review of Akh: Une Notion Religieuse Dans L’Égypte Pharaonique by Gertie Englund. J Am Res Cent Egypt. 1982;19:145–8.

35. Friedman F. Akh. In: Redford DB, editor. Oxford Encyclopedia of Ancient Egypt I.Oxford: Oxford University Press; 2001.

36. Smith M. Traversing eternity: texts for the afterlife from Ptolemaic and Roman Egypt. Oxford:Oxford University Press; 2009.

37. Janák J. Staroegyptské náboženství II. Život a úděl člověka. Praha: Oikoymenh; 012.

38. Assmann J. Tod und Jenseits im alten Ägypten. München: C.H. Beck; 2001.

39. Janák J. Northern Bald Ibis (Akh-Bird). In: UCLA Encyclopedia of Egyptology [Internet]. Los Angeles: University of California in Los Angeles; 2013. Available from: http://digital2.library.ucla.edu/viewItem.do?ark=21198/zz002dwqt8

40. Leprohon RJ. Gatekeepers of This and the Other World. J Soc Study Egypt Antiq. 1994;24:77– 91.

41. Jansen-Winkler K. “Horizont” und “Verklärheit”: Zur Bedeutung der Wurzel Ax. Stud zur Altägyptischen Kult. 1996;23:201–15.

42. Allen JP. The Cosmology of the Pyramid Texts. In: Religion and Philosophy in Ancient Egypt, Yale Egyptological Studies 3. New Haven: Yale Universitxy Press; 1989. p. 1–29.

43. Allen JP. Reading a Pyramid. In: C. Berger, editor. Homages à Jean Leclant Volume 1: Études pharaoniques. Cairo: Orientale, Institut Français d’Archéologie; 1994.

44. Hays HM. Unreading the Pyramids. Le Bull l’Institut français d’archéologie Orient. 2009;195–220.

45. Demarée RJ. The Ax iqr n Ra-stelae on Ancestor Worship in Ancient Egypt. Leiden; 1983.

46. Boehm C, Pegoraro K. Der Waldrapp: Ein Glatzkopf in Turbulenzen. Neue Brehm. Hohenwarsleben: Westarp Verlag; 2011.

47. Fritz J, Unsoeld M. Aufwind für den Waldrapp: Von der Wiederansiedlung eines europäischen Zugvogels. Jahrb des Vereins zum Schutz der Bergwelt. 2013;78:121–38.

48. Boessneck J. Die Tierwelt des Alten Ägypten. Munich: C.H. Beck; 1988.

49. Janák J. A question of size: A remark on early attestations of the ba hieroglyph. Stud zur Altägyptischen Kult. 2011;143–53.

50. Dunham D. An Egyptian diadem of the Old Kingdom. Bull Museum Fine Arts. 1946;23–9.

51. Staehelin E. Untersuchungen zur ägyptischen Tracht im Alten Reich (doctoral thesis). Hessen, Berlin; 1966.

52. Kees H. Der Opfertanz des ägyptischen Königs. München; 1912.

53. Decker M, Herb W. Bildatlas zum Sport im alten Ägypten. Corpus der bildlichen Quellen zu Leibesübungen, Spiel, Jagd, Tanz und verwandten Themen. Leiden, New York & Cologne: Brill; 1994.

54. Zoufal K, Fritz J, Bichler M, Kirbauer M, Markut T, Meran I, et al. Feeding ecology of the Northern Bald Ibis in different habitat types: An experimental field study with handraised individuals. Report of the 2nd IAGNBI Meeting. 2007.

55. Fritz J, Wirtz S, Unsoeld M. Aspekte der Nahrungsökologie und Genetik des Waldrapps: Reply zu Bauer & etal. (2016) Vogelneozoen in Deutschland - Revision der nationalen Statuseinstufungen. Vogelwarte. 2017;55:141–5.

56. Serra G, Abdallah MS, al Qaim G. Feeding ecology and behaviour of the last known survivi8ng Northern Bald Ibises, Gerontius eremita, at theier breeing quarter in Syria. Zool Middle East. 2008;43:55–68.

57. Serra G, Bruschini C, Peske L, Kubsa A, Wondafrash M, Lindsell JA. An assessment of ecological conditions and threats at the Ethiopian wintering site of the last known eastern colony of Critically Endangered Northern Bald Ibis Geronticus eremita. Bird Conserv Int. 2013;150(4).

58. Bowden CGR, Smith KW, Bekkay M El, Oubrou W, Aghnaj A, Jimenez-Armesto M. Contribution of research to conservation action for the Northern Bald Ibis Geronticus eremita in Morocco. Bird Conserv Int. 2008;18:74–90.

59. Serra G, Peske L, Abdallah MS, al Qaim G, Kanani A. Breeding ecology and behaviour of the last wild oriental Northern Bald Ibises (Geronticus eremita) in Syria. J Ornithol. 2009;150(4):769– 82.

60. Schenker A, Cahenzli F, Gutbrod KG, Thevenot M, Erhardt A. The Northern Bald Ibis Geronticus eremita in Morocco since 1900: Analysis of ecological requirements. Bird Conserv Int [Internet]. 2019;1–22. Available from: https://www.cambridge.org/core/product/identifier/S0959270919000170/type/journal_article

61. Yeniyurt C. Conservation of the Northern Bald Ibis, Birecik, Turkey 2013-2016. Report of 4th IAGNBI Meeting Seekirchen (Ed. C Boehm & C Bowden). 2016.

62. Smith KW, Aghnaj A, El Bekkay M, Oubrou W, Ribi M, Armesto MJ, et al. The provision of supplementary fresh water improves the breeding success of the globally threatened Northern Bald Ibis Geronticus eremita. Ibis (Lond 1859). 2008;150(4):728–34.

63. Sorato E, Kotrschal K. Hormonal and behavioural symmetries between the sexes in the Northern bald ibis. Gen Comp Endocrinol. 2006;146(3):265–74.

64. Voelkl B, Fritz J. Relation between travel strategy and social organization of migrating birds with special consideration of formation flight in the northern bald ibis. Philos Trans R Soc B Biol Sci [Internet]. 2017;372(1727):20160235. Available from: http://rstb.royalsocietypublishing.org/lookup/doi/10.1098/rstb.2016.0235

65. Pegoraro K, Föger M. Die ‘Chrup’-Rufe des Waldrapps Geronticus eremita: Ihre verschiedenen Funktionen in einem komplexen Sozialsystem. J Ornithol. 1995;136:243–52.

66. Szipl G, Boeckle M, Werner SAB, Kotrschal K. Mate recognition and expression of affective state in croop calls of Northern Bald Ibis (Geronticus eremita). PLoS One. 2014;

67. Pegoraro K, Föger M. Individuality in the Northern Bald Ibis or Waldrapp Ibis Geronticus eremita – key features for a complex social system. Acrocephalus. 2001;22:73–9.

68. Sorato E, Kotrschal K. Skin ornaments reflect social status and immunocompetence in male and female Northern Bald Ibises (Geronticus eremita). unpublished data.

69. Galván I, Palacios D, Negro JJ. The bare head of the Northern bald ibis (Geronticus eremita) fulfills a thermoregulatory function. Front Zool. 2017;14(1):1–9.

70. Unsoeld M, Melzer R. Sunning behaviour in Ibises (Threskiornithidae): Observations on four species at Tierpark Hellabrunn, Munich Markus. Der Zooloigsche Garten. 2010;79(2–3):89–104.

71. Fritz J, Kramer R, Hoffmann W, Trobe D, Unsoeld M. Back into the wild: establishing a migratory Northern bald ibis Geronticus eremita population in Europe. Int Zoo Yearb. 2017;51(1):107–23.

72. Hegemann A, Fudickar AM, Nilsson J-Å. A physiological perspective on the ecology and evolution of partial migration. J Ornithol [Internet]. 2019;(August 2018). Available from: http://link.springer.com/10.1007/s10336-019-01648-9

73. Fritz J, Feurle A, Kotrschal K. Physiological regulation of bird migration: a study with northern bald ibises undergoing human-led autumnal migration. J Ornithol. 2016;147(5):168.

74. Bairlein F, Fritz J, Scope A, Schwendenwein I, Stanclova G, Van Dijk G, et al. Energy expenditure and metabolic changes of free-flying migrating northern bald ibis. PLoS One. 2015;10(9).

75. Fritz J. Ultraleichtflieger weisen den Weg – Der Waldrapp in den Alpen. Der Falke. 2010;57:95– 105.

76. Sperger C, Heller A, Voelkl B, Fritz J. Flight Strategies of Migrating Northern Bald Ibises – Analysis of GPS Data During Human-led Migration Flights. Agit ‒ J für Angew Geoinformatik. 2017;3:62–72.

77. Portugal SJ, Hubel TY, Fritz J, Heese S, Trobe D, Voelkl B, et al. Upwash exploitation and downwash avoidance by flap phasing in ibis formation flight. Nature. 2014;505(7483):399–402.

78. Voelkl B, Portugal SJ, Unsoeld M, Usherwood JR, Wilson AM, Fritz J. Matching times of leading and following suggest cooperation through direct reciprocity during V-formation flight in ibis. Proc Natl Acad Sci [Internet]. 2015;112(7):2115–20. Available from: http://www.pnas.org/lookup/doi/10.1073/pnas.1413589112

79. Voelkl B, Fritz J. Relation between travel strategy and social organization of migrating birds with special consideration of formation flight in the northern bald ibis. Philos Trans R Soc B Biol Sci. 2017;372(1727).

80. Žabkar L. A Study of the Ba Concept in Ancient Egyptian Texts. Chicago: The Universtiy of Chicago Press; 1968.

81. Aourir M, Bousadik H, El Bekkay M, Oubrou W, Znari M, Qninba A. New Breeding Sites of the Critically Endangered Northern Bald Ibis Geronticus Eremita on the Moroccan Atlantic Coast. Int J Avian Wildl Biol. 2017;2(3):1–4.

82. Touti J, Oumellouk F, Bowden CGR, Kirkwood JK, Smith KW. Mortality incident in northern bald ibis Geronticus eremita in Morocco in May 1996. Oryx. 1999;33(2):160–7.

83. Bowden CGR (Compiler). International Single Species Action Plan for the Conservation of the Northern Bald Ibis (Geronticus eremita). Vol. 55. Bonn, Germany; 2015.

84. Fritz J, Unsoeld M. Internationaler Artenschutz im Kontext der IUCN Reintroduction Guidelines: Argumente zur Wiederansiedlung des Waldrapps Geronticus Eremita in Europa. Vogelwarte. 2015;53(2):157–68.

85. Drenske S, Radchuk V, Scherer C, Esterer C, Kowarik I, Fritz J, et al. From reintroduction to rewilding: Northern bald ibis (Geronticus eremita) still need management interventions for critical survival. in prep.

86. Ikram S. Divine creatures: Animal mummies in ancient Egypt. Cairo: American University in Cairo Press; 2005.

87. Houlihan P. The birds of ancient Egypt. Cairo: American University in Cairo Press; 1988.

88. Bárta M. Kolaps Staré říše, éry stavitelů pyramid?In: P. POKORNÁ; M. BÁRTA, editor. Něco překrásného se končí Kolapsy v přírodě a společnosti. Praha: Dokořán; 2008. p. 121–44.

89. Dalfes HN, Kukla G, Weiss H. Third millennium BC climate change and old world collapse. Berlin/New York: Springer; 1997.

90. Issar AS, Zohar M. limate change: environment and civilization in the Middle East. Berlin/New York: Springer; 2004.

91. Janák J. Saddle-billed Stork (ba-bird). In: UCLA Encyclopedia of Egyptology [Internet]. Los Angeles: University of California in Los Angeles; 2014. Available from: http://digital2.library.ucla.edu/viewItem.do?ark=21198/zz002hp1vp

92. Schilling J, Freier KP, Hertig E, Scheffran J. Climate change, vulnerability and adaptation in North Africa with focus on Morocco. Agric Ecosyst Environ [Internet]. 2012;156(January 2018):12–26. Available from: http://dx.doi.org/10.1016/j.agee.2012.04.021

93. Visser ME, Perdeck AC, van Balen JH, Both C. Climate change leads to decreasing bird migration distances. Glob Chang Biol. 2009;15(8):1859–65.

94. Radchuk V, Reed T, Teplitsky C, van de Pol M, Charmantier A, Hassall C, et al. Adaptive responses of animals to climate change are most likely insufficient. Nat Commun [Internet]. 2019;10(1):3109. Available from: http://www.nature.com/articles/s41467-019-10924-4

95. Boehm C, Fritz J, Asmus J. Koordination und Kooperation von Zoo-und Freilandarbeit bis zur Wiederansiedlung: vier Fallbeispiele. In: W. Lantermann, Asmus J, editors. Wildvogelhaltung. Springer-Verlag GmbH; 2021.

96. Lelieveld J, Hadjinicolaou P, Kostopoulou E, Chenoweth J, El Maayar M, Giannakopoulos C, et al. Climate change and impacts in the Eastern Mediterranean and the Middle East. Clim Change. 2012;114(3–4):667–87.

